# The Allen Ancient DNA Resource (AADR): A curated compendium of ancient human genomes

**DOI:** 10.1101/2023.04.06.535797

**Authors:** Swapan Mallick, Adam Micco, Matthew Mah, Harald Ringbauer, Iosif Lazaridis, Iñigo Olalde, Nick Patterson, David Reich

**Affiliations:** Department of Genetics, Harvard Medical School, Boston, MA 02115, USA; Broad Institute of MIT and Harvard, Cambridge, MA 02142, USA; Howard Hughes Medical Institute, Boston, MA 02115, USA; Department of Human Evolutionary Biology, Harvard University, Cambridge, MA 02138, USA; Max Planck Institute for Evolutionary Anthropology, Leipzig 04103, Germany; BIOMICs Research Group, University of the Basque Country, 01006 Vitoria-Gasteiz, Spain

## Abstract

More than two hundred papers have reported genome-wide data from ancient humans. While the raw data for the vast majority are fully publicly available testifying to the commitment of the paleogenomics community to open data, formats for both raw data and meta-data differ. There is thus a need for uniform curation and a centralized, version-controlled compendium that researchers can download, analyze, and reference. Since 2019, we have been maintaining the Allen Ancient DNA Resource (AADR), which aims to provide an up-to-date, curated version of the world’s published ancient human DNA data, represented at more than a million single nucleotide polymorphisms (SNPs) at which almost all ancient individuals have been assayed. The AADR has gone through six public releases since it first was made available and crossed the threshold of >10,000 ancient individuals with genome-wide data at the end of 2022. This note is intended as a citable description of the AADR.

---

The first genome-wide ancient DNA data were published in 2010 [1-3]. However, it was only in 2015 with the advent of large-scale studies of Holocene genomes, in-solution enrichment of ancient DNA libraries for targeted single nucleotide polymorphisms (SNPs) [4-6] and the introduction of automated protocols and liquid handling robots for processing of ancient DNA libraries [7, 8], that the number of individuals with genome-wide data began to increase rapidly. Between 2010 and 2014, data from an average of about 10 individuals with genome-wide data were published each year. Between 2015 and 2017, the numbers increased to about 200 annually. Since 2018, data from thousands of individuals have been published every year (Figure 1). Ancient DNA data to date has been concentrated in western Eurasia, but an increasingly large fraction of the data derive from other regions of the world (Figure 2).

**Figure 1:**
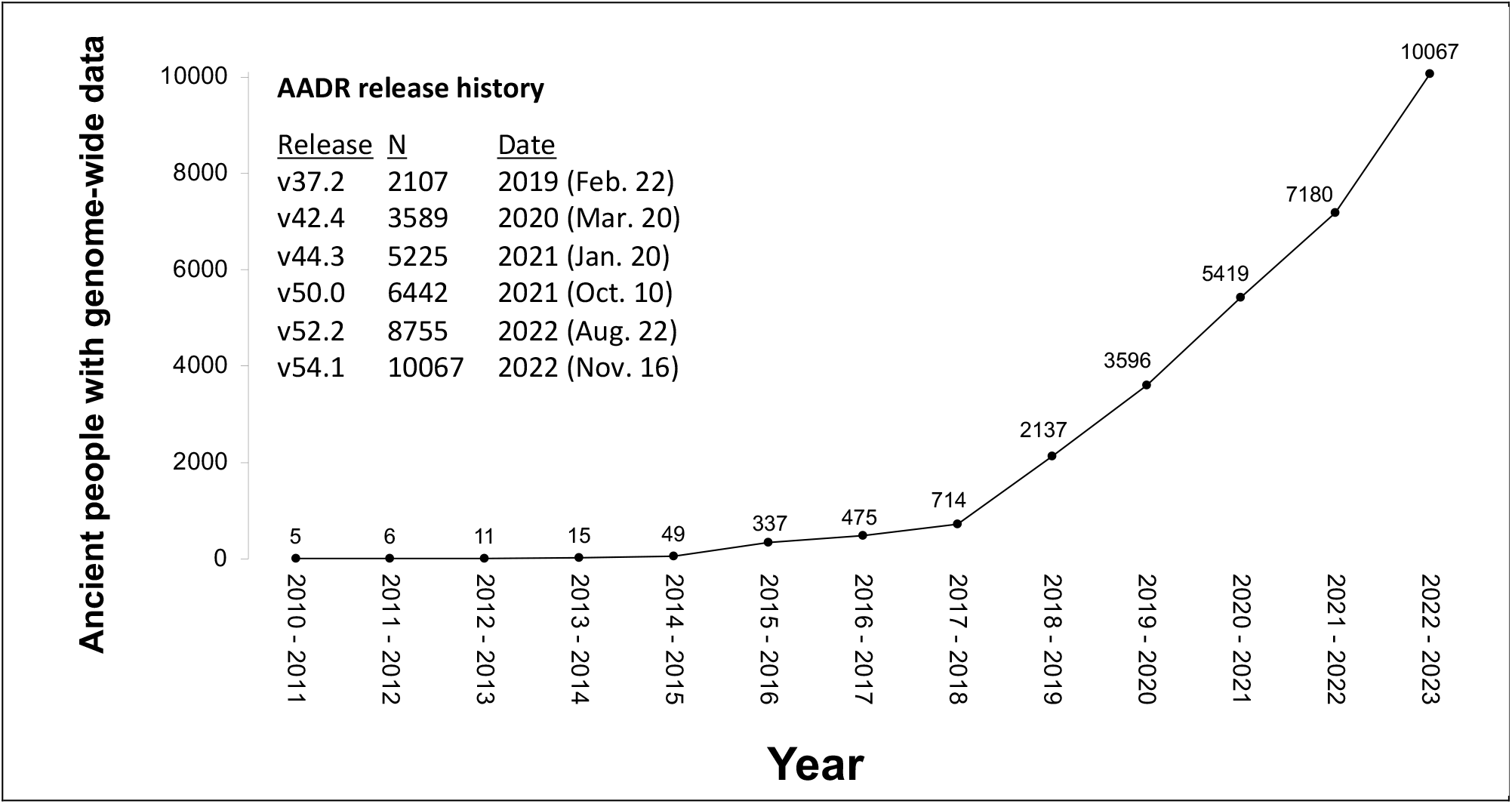
Growth in the number of humans with published genome-wide ancient DNA data Ancient people with genome-wide data.

**Figure 2:**
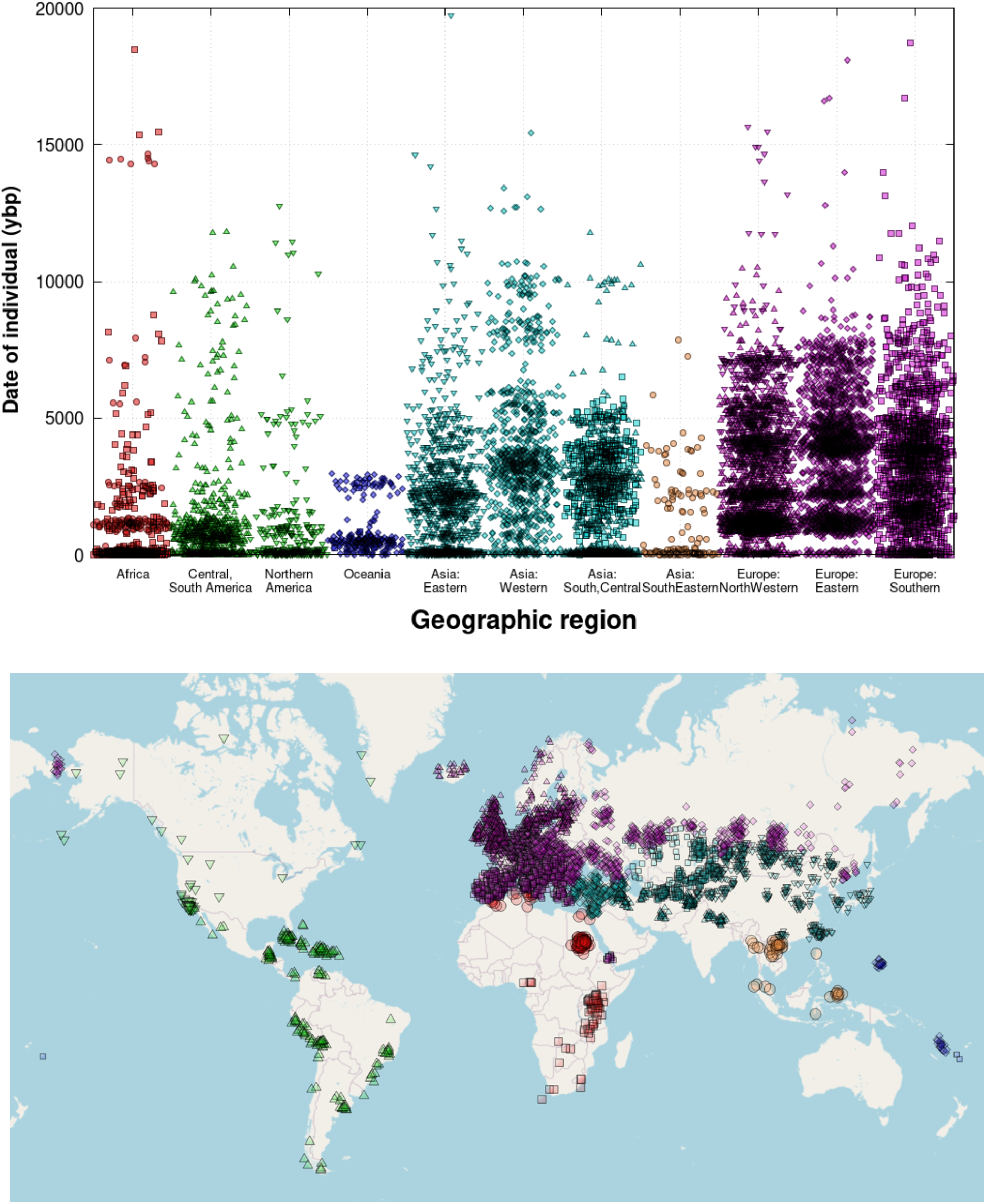
Geographical and temporal distribution of individuals in the v54.1 AADR. (released November 2022). In the upper panel, we do not show 17 individuals older than 20,000 years.

A challenge in analyzing ancient DNA is that individuals are distributed over many independent studies. Thus, while raw sequence data for more than 99% of individuals are fully available in public repositories [9] the uploaded data exist in diverse formats, as does the meta-data such as archaeological, chronological, and geographic information. Some resources exist which consolidate subsets of publicly available ancient DNA data, including a Y-chromosome database with assembled data from nearly two thousand ancient Eurasian Y-chromosome haplogroups [10], a mitochondrial DNA database with more than two thousand individuals [11], and the Online Ancient Genome Repository [12] which copies publicly available data and encapsulates each dataset into an archived tar file. However, none of these provide a curated dataset that attempts to include all published data in a single easily co-analyzable format.

The aim of the Allen Ancient DNA Resource (AADR) is to maintain an up-to-date representation of the world’s published human genome-wide ancient DNA data, in a format that allows meaningful co-analysis of data generated across different studies. Work on harmonization is important, as there can be considerable heterogeneity in data produced in different studies, which has potential to confound analysis. For example, there are major differences in library preparation protocols, with some library preparations using treatment with the enzyme uracil DNA glycosylase to the reduce errors due to post-mortem cytosine deamination which primarily affects the ends of molecules, and some not doing this because the treatment reduces library complexity [7]. Another heterogeneity is associated with whether data are produced by in-solution enrichment of ancient DNA libraries at a targeted set of more than a million SNPs (about 70% of individuals with published data) or shotgun sequencing (almost all of the remaining individuals). While in-solution enrichment has the advantage of increasing the rate of converting sampled individuals into working genome-wide data and decreasing the cost of analysis per individuals, it can produce biases toward one allele or the other at analyzed SNPs that can make it challenging to co-analyze data produced with this method and data produced using direct shotgun sequencing [13]. There are also differences around computational processing of raw sequencing data, which can produce batch effects across studies [14, 15].

To bring data generated outside our own laboratory into the AADR, we usually start with available sequences from a public repository, most often the European Nucleotide Archive (https://www.ebi.ac.uk/ena), following accession numbers given in the published papers. In some cases we start with alternatively formatted versions that we request directly from the authors. For data generated in our laboratory, we start from our own raw sequence files, which are the basis for data uploaded to established public repositories.

The raw data we analyze come in diverse formats, usually *fastq* files (for raw sequence data) or *bams* (for either unaligned reads or reads aligned to a reference genome) [16]. A challenge is that there can be considerable variation in *fastq* and *bam* files, reflecting the formatting, filtering and processing choices made by researchers in generating data. This includes:

(a) Base calls and associated quality scores in raw sequences are often modified by the researchers who generated the data. One common modification is to recalibrate base quality scores [17]. Another modification is to filter out the ends of sequences, either by masking terminal bases in the sequences that are uploaded and marking them as “N”, or clipping them (removing) altogether [18]. Such filtering aims to greatly reduce error rates associated with cytosine deamination typical of ancient DNA data. However, it also means that users cannot make choices about whether to use the valuable data that has been masked and clipped (such as sites unaffected by deamination). In addition, it has the effect of making it difficult to identify damaged molecules which are a strong indicator that those molecules indeed are ancient and not derived from some potential contaminating modern human source.
(b) Sequences may be aligned to different human reference genomes, typically hg19, hs37d5, or hg20, each with their own unique coordinate systems. To build a homogeneous dataset, we therefore have to map to a unified coordinate system, currently based on hg19 [19-21]. A further challenge is that chromosomes may have inconsistent naming conventions, (for example ‘chr1’ v. ‘1’, or ‘chrMT’ v. ‘MT’ v. ‘chrM’), or the sorting order of chromosomes can differ. This results in practical difficulties in merging datasets.
(c) Data may be deposited either (i) by library, or (ii) by-individual with multiple libraries in a single file. If data are deposited by library, then it may be necessary to identify and perform a merging step. There are pitfalls that arise in such merging, as in some cases “readgroup” names (a tag which groups reads together) are the same across individuals, and so joint processing of many individuals can inadvertently lead to in-silico contamination.

For the AADR, we manually process the datasets from each individual, tailoring each according to its characteristics. We then create a *bam* file aligned to the hg19 genome reference sequence. The bam files used to generate AADR constitute tens of terabytes in size altogether. For the AADR, we process these bams to produce “pseudohaploid” genotypes at a set of more than a million SNPs that have been assayed for nearly all published individuals with ancient DNA data; by “pseudohaploid”, we mean that we represent the individual by a randomly sampled sequence at each analyzed position. For individuals where coverage is sufficient to allow full genotyping, we also release diploid genotypes [22, 23]. We finally manually curate the genotypes to check that samples have the expected population genetic properties.

AADR consists of three standard files in EIGENSTRAT format (.*ind*, .*snp*, and .*geno*). We also include an annotation file that is rich in meta-information for the dataset (.*anno*). The .*anno* file includes meta-data manually extracted from the papers reporting the data, in some cases supplemented by information that appeared later or reflect clarifications from original sources. For archaeological information, we attempt to provide:

- Skeletal codes and grave numbers and sometimes other identifiers always also including the code used for genetic analysis.
- Latitude and longitude.
- Location information, with a separate column for “Political entity” such as country, and locality information with largest region first, and more specific information thereafter.
- Chronological information in a standard format. When a radiocarbon date is available, we include the laboratory number and calibrated 95.4% confidence interval obtained in OxCal v4.4.2 using either the IntCal20 or SHCal20 calibration curve (if we make an alternative choice, it is explicitly explained in a Methods column). We also report the posterior mean and standard deviation of the calibrated radiocarbon date. When no radiocarbon date is available, we present a date uncertainty range based on archaeological context, usually rounded to the nearest 50 or 100 years, and quote the mean and standard deviation assuming a uniform distribution over the range.
- We include a group name for the individual, using a naming convention that aims to be systematic [24]. The group name may additionally include a suffix that mark individuals such as potentially contaminated (“_contam”), or as a population genetic outlier (“_o”), or as having relatively little data (low coverage – “lc”). Data from individuals generated using shotgun sequencing methods have a suffix “.SG” (for pseudohaploid representations) or “.DG” (for diploid representations).
- We include an estimate of the age of the individual at their death based on physical anthropology when we are able to obtain it.
- We include many metrics computed on the genetic data, including not just amount of data (average coverage assayed at the subset of 1.15 million autosomal sites targeted in the 1.2 million SNP enrichment assays), but also molecular sex determination, cytosine-to-thymine rate in the final nucleotide [25], fraction of the genome in multi-megabase runs of homozygosity [26], identification of close relatives in the dataset (in a dedicated “family information” column), and estimates of contamination [27, 28].
- When data from an individual have been published in multiple papers using the same methodology such as in-solution enrichment, the AADR typically includes only the best quality version which is usually the latest one (for such individuals, the “publication” columns in the .*anno* file note the publication that first reported the data as well as the publication that report the version that is actually included within the AADR). For some individuals, we include multiple representations of data, for example data from shotgun sequencing, data from in-solution enrichment, data restricted to UDG-treated libraries, or data restricted to sequences showing characteristic ancient DNA damage to reduce the possible impact of contaminating sequences (“_d” suffix). The different versions have unique “Genetic IDs” but the same “Master ID” (which uniquely identifies an individuals).

Because we are trying to keep AADR current, we err on the side of inclusivity, and thus bring data into the dataset even when meta-information and metrics are incomplete. Each AADR release updates the meta-information and identifiers as appropriate. We rely on ongoing curation of the dataset as well as feedback from the user community which we invite through communication with the corresponding authors, to identify individuals with erroneous meta-information or corrupted genetic data, which we then seek to correct in subsequent releases.

To increase the usefulness of the AADR, we have added in data from diverse modern humans, including shotgun sequencing data from sets of individuals included within the 1000 Genomes Project [29], the Simons Genome Diversity Project [30], and the Human Genome Diversity Project [31]. To integrate these data, we have had to address challenges of different reference genomes (for example transforming from hg20 to hg19 coordinates). There are 6399 modern individuals with shotgun data in the v54.1 release.

We have also integrated a dataset of 4114 modern individuals genotyped on the Affymetrix Human Origins array at approximately ∼600,000 SNPs [32]. This is a sufficiently valuable dataset that the AADR provides two releases: one on all ∼1.2 million targets (excluding the Human Origins data), and one on just the Human Origins targets.

Since the v52.2 release, we have also maintained a mitochondrial repository, which includes mitochondrial genomes for 4122 ancient individuals in the AADR.

The AADR is now on its sixth public release (v54.1). It is freely available from: https://reich.hms.harvard.edu/allen-ancient-dna-resource-aadr-downloadable-genotypes-present-day-and-ancient-dna-data, with version-controlled releases including permanent digital object identifiers (doi’s) for each release available from the Harvard Dataverse (accession: https://doi.org/10.7910/DVN/FFIDCW). Researchers who use the curated dataset from the AADR as the basis for analyses should cite this paper and the version of the AADR they downloaded, including a reference to that version’s doi. Citing the AADR paper is not a substitute for citing the original publications that produced data, which should be specifically referenced in each publication.

## Acknowledgements

We are grateful to Rebecca Bernardos, Aisling Kearns, Nadin Rohland, Arie Shaus, Katie Mika and many in the user community for help in curating and improving the AADR. Construction and maintenance of the AADR was supported by NIH grant HG012287, by the Allen Discovery Center program, a Paul G. Allen Frontiers Group advised program of the Paul G. Allen Family Foundation, by John Templeton Foundation grant 61220, and by the Howard Hughes Medical Institute. This article is subject to HHMI’s Open Access to Publications policy. HHMI lab heads have previously granted a nonexclusive CC BY 4.0 license to the public and a sublicensable license to HHMI in their research articles. Pursuant to those licenses, the author-accepted manuscript of this article can be made freely available under a CC BY 4.0 license immediately upon publication.

## Notes

https://reich.hms.harvard.edu/allen-ancient-dna-resource-aadr-downloadable-genotypes-present-day-and-ancient-dna-data

https://doi.org/10.7910/DVN/FFIDCW

### Competing Interest Statement

The authors have declared no competing interest.

